# Poloxamer dilution as an on-demand alternative to agar dilution-based antimicrobial susceptibility testing

**DOI:** 10.64898/2025.12.09.693179

**Authors:** Matthew T.J. Uy, Andrea Kirmaier, Lindsey M. Rudtner, Aidan Pine, James E. Kirby

## Abstract

Agar dilution is a reference susceptibility testing method uniquely or preferentially recommended for certain antimicrobials. However, the effort required to pour individual agar plates spanning a doubling dilution range precludes its practical implementation in hospital clinical laboratories. Here, we describe an on-demand replacement for agar dilution (AD), specifically substituting poloxamer 407 (also known as Pluronic F-127) for Bacto Agar as the solidifying agent. Notably, 20% poloxamer 407 solutions (e.g., with Mueller-Hinton broth) remain liquid at refrigerated temperatures, but solidify upon warming, enabling facile set-up of poloxamer dilution testing (PD) in Petri dish or microwell format. For fosfomycin susceptibility testing, PD and reference agar dilution (AD) showed excellent categorical (CA) and essential (EA) agreement for *E. coli* (100% and 87%, respectively, n=31). For other *Enterobacterales,* excluding *Klebsiella* spp; CA and EA were both 82% (n=17, respectively). For *P. aeruginosa*, CA and EA were 60% and 100% (n=10), respectively, with the lower CA reflecting the large number of strains tested with MICs near categorical breakpoints. There were no very major errors, while major errors were only observed for *Klebsiella* spp. Additionally, PD substantially reduced the number of skipped dilutions 6-fold for *E. coli* (P < 0.0001) and inhibited swarming of *Proteus* species. We conclude that PD and AD, an imperfect gold standard, have essentially equivalent practical performance and that PD can therefore serve as an on-demand alternative testing methodology in clinical laboratories for fosfomycin testing of gram-negative pathogens. A broader exploration of PD’s utility is thus warranted.

**Importance:** Accurate antibiotic susceptibility testing is essential for guiding treatment of bacterial infections. For the antibiotic fosfomycin, used to treat *Escherichia coli* urinary tract infections, the most reliable testing method requires solid media prepared by hand for each antibiotic concentration, which is too time consuming for most clinical laboratories to perform. Our study shows that replacing agar with an alternative temperature-sensitive gelling agent called poloxamer enables laboratories to prepare solid test plates rapidly without special equipment. This approach, which is essentially identical to traditional agar dilution, provides a practical alternative that can be implemented by hospital clinical microbiology laboratories for accurate fosfomycin testing near the point of patient care. This strategy may also be applicable to other drugs for which agar dilution is the preferred testing method, supporting expedited testing to inform treatment decisions for bacterial infections.

## Introduction

Antimicrobial resistance is emerging as a significant concern for the treatment of bacterial infections. To identify appropriate therapies for bacterial infections, clinical laboratories must therefore routinely conduct antimicrobial susceptibility testing (AST) on patient isolates. Manually prepared broth or agar antimicrobial susceptibility testing panels are considered the reference gold standard for the determination of minimal inhibitory concentrations values (MIC). In broth microdilution, the MIC is defined as the lowest concentration of an antimicrobial that visibly inhibits bacterial growth after a specified incubation period (1–3). For agar dilution, the MIC corresponds to the lowest agar plate concentration, in a twofold dilution series, that shows no visible growth of the test isolate. MICs are categorized are susceptible, susceptible dose-dependent, intermediate, or resistant based on CLSI, FDA, or EUCAST standards to guide antibiotic therapy selection.

Existing reference AST methods are labor-intensive and technically complex, precluding their use in most hospital clinical laboratories near the site of patient care. Therefore, standard hospital-based clinical laboratory AST is generally performed using automated commercial platforms based on variants of broth dilution testing or alternatively by disk diffusion or gradient diffusion methods (4). However, these methods may not be appropriate for testing diffusion-limited drugs.

For certain drug-organism combinations, agar dilution is either the only or preferred CLSI testing method (1–3). A prominent example is fosfomycin, an antimicrobial commonly prescribed for treatment of uncomplicated urinary tract infections (uUTIs) for which agar dilution serves as the reference method. Broth microdilution and gradient diffusion methods exhibit suboptimal accuracy and reproducibility. However, agar dilution itself is associated with a high frequency of skipped dilutions (the equivalent of “skipped wells” in broth microdilution testing) (5, 6), which precludes the reliable use of single-concentration agar breakpoint screening plates, as such irregularities could yield unrecognized errors.

CLSI fosfomycin disk diffusion breakpoints are available for *E. coli* and *E. faecalis*, but not for other organisms, because the drug’s FDA approval is limited to treatment of uUTI caused by these pathogens. Moreover, disk diffusion-based methods correlate poorly with agar dilution testing outside these organism groups (7). Nevertheless, because fosfomycin may be the only oral agent with potential activity against certain gram-negative urinary pathogens, clinical laboratories are frequently asked to extrapolate *E. coli* disk diffusion breakpoints to other organisms.

It would be ideal if accurate fosfomycin MIC results for *E. coli* and other organisms could be generated near the site of patient care. This need arises because reference laboratory testing results are necessarily delayed, and reference laboratories also generally do not perform MIC testing in the absence of species CLSI breakpoints.

We have identified a potential way to address this issue based on technology developed in prior work. Previously, our group developed a rapid microscopy-based antimicrobial susceptibility testing (MAST) platform that met FDA accuracy criteria within 4 hours of organism inoculation (8). As part of this work, we found that organisms could reliably grow on an optically clear Mueller–Hinton medium solidified with Poloxamer 407, which fully supported the growth of a wide range of gram-negative and gram-positive pathogens. Poloxamer 407 possesses a unique and beneficial physical property in solution: it remains liquid when refrigerated but transitions to a solid gel at higher temperatures (9–11). We therefore hypothesized that a cold, sterile liquid poloxamer stock solution could substitute for standard Bacto agar as the solidification reagent, providing a flexible and readily implemented alternative to agar dilution.

The potential advantages of such a solidification agent are numerous. Poloxamer media aliquots can be sterilized by autoclaving and stored at 4 °C until use, then gently mixed with doubling dilutions of antibiotics, solidified at room temperature in less than 30 minutes, and subsequently inoculated and read using standard agar dilution protocols. This approach could therefore enable an agar dilution-equivalent method that can be prepared easily and on demand, eliminating the need for the repetitive, time-consuming process of autoclaving media, holding molten agar at a narrow temperature range, and manually incorporating doubling dilutions of antibiotics.

We therefore sought to validate poloxamer dilution technology compared with reference agar dilution testing, using fosfomycin antimicrobial susceptibility testing as a representative use case.

## Materials and Methods

### Bacterial Strains

*Escherichia coli* ATCC 25922 and *Enterococcus faecalis* ATCC 29212 were obtained from the ATCC. *E. coli* AR 346 and AR-549 were *fosA* isolates from the FDA-CDC AR-Bank collection (12). Additional de-identified, colony-purified clinical isolates were retrieved from the Beth Israel Deaconess Medical Center clinical microbiology laboratory as listed in **Table S1** under an IRB-approved protocol.

## Medial Preparation

The following reagents were used for media preparation: Pluronic F-127 (Sigma-Aldrich, P2443-250G, lot BCBV1572), non–cation-adjusted Mueller-Hinton broth (BD Difco, 275730, lot 3150802), Bacto agar (BD DF0140-15-4), D-glucose 6-phosphate potassium salt (Sigma-Aldrich, G6526-1G, lot SLBH9805V), and fosfomycin disodium salt (Sigma-Aldrich, P5396, lot 125M4169).

## Base Media Preparation

### Mueller–Hinton–Poloxamer (MH-P) Base

Poloxamer 407 was dissolved in deionized water to a final concentration of 25% (w/v) by stirring overnight at 4 °C. One-fifth volume of 5× Mueller–Hinton broth (10.5% w/v) was then added to yield a final mixture containing 20% poloxamer 407 and 1× Mueller–Hinton broth. Filter-sterilized D-glucose-6-phosphate potassium salt was added to a final concentration of 25 µg/mL. The solution was autoclaved, cooled, and stored at 4 °C in liquid form until use.

### Agar Dilution (AD) Base

Mueller–Hinton agar was prepared by combining Mueller–Hinton broth with 1.5% (w/v) Bacto agar, autoclaving, and maintaining the molten medium at ∼50 °C in a water bath. Filter-sterilized D-glucose-6-phosphate potassium salt was then added to a final concentration of 25 µg/mL.

### Agar Dilution (AD) Plates

Doubling dilutions of fosfomycin (512 to 0.25 µg/mL) were prepared by adding appropriate volumes of filter-sterilized fosfomycin stock to aliquots of AD base medium. Plates (15 mL per plate) were poured, allowed to solidify at room temperature, and used the following day.

### Poloxamer Dilution (PD) Plates

For PD plates, filter-sterilized fosfomycin stock was added to pre-made MH-P base aliquots stored at 4 °C, gently mixed by swirling, and dispensed into plates (15 mL per plate). Plates were allowed to solidify at room temperature prior to use.

### Accuracy and Precision Analysis

Fosfomycin susceptibility testing of all controls and clinical isolates (n=82) was performed in parallel using the PD and AD method reference method. For all clinical isolates examined, susceptibility testing was repeated three times, each biological replicate performed on a separate day for both methods in parallel. Prior to each day of testing, isolates were re-streaked on nonselective blood agar plates and incubated at 37°C for 18 to 24 hours. Single colonies from isolates were suspended in sterile Mueller Hinton broth to 0.5 McFarland using a BioMérieux DensiCHEK Plus instrument. 1:10 dilutions of the suspensions were loaded into inoculum wells of a sterilized Steers Replicator, allowing delivery of 2 µL or 10^4^ CFU per spot onto PD and AD doubling plates in parallel in 35 isolate batches. Plates were incubated at 37°C for 16 to 20 hours and visually scored to determine the MIC of each strain, defined as the lowest antimicrobial concentration at which bacterial growth was completely inhibited. *Escherichia coli* ATCC 25922 and *Enterococcus faecalis* ATCC 29212 were also tested in parallel with each experiment and were within acceptable quality control ranges for both PD and AD for all experiments. Two fosfomycin-resistant, *fosA* AR-Bank *E. coli* isolates and a fosfomycin susceptible *E. coli* clinical isolate were also included on each day of testing.

For method comparison, a single consensus MIC was assigned to each isolate for each method. If at least two of the three replicates agreed, the modal MIC was used. If all replicate MICs differed or only two valid replicates were available (e.g., due to skipped dilutions), and they disagreed, the median doubling dilution (rounded to the nearest dilution when between tested values) was assigned. Skipped dilutions were handled per CLSI M07 guidelines with the MIC read above the highest concentration with observed growth. Isolate readings with more than one consecutive or non-consecutive skipped dilutions were considered invalid. These consensus MICs were then compared between PD and AD to calculate essential agreement (EA; ±1 doubling dilution) and categorical agreement according to established guidelines (16). For all agreement calculations, AD results were used as the reference standard. Results were also considered in EA if both PD and AD yielded the same off-scale results (i.e. >512). Categorical Agreement (CA) was established if both PD and AD yielded the same categorical result. Very major errors (VME) were defined as PD susceptible and AD resistant; major errors (ME) as PD resistant and AD susceptible; and minor errors (MinE) as one result intermediate and the other susceptible or resistant. Wilson Score confidence intervals were calculated using RStudio.

Precision essential agreement (PEA) analysis for the entire data set was determined by assessing consistency among triplicate MIC determinations for each isolate. PEA was defined as all three replicates yielding MICs within ±1 doubling dilution of each other. Precision categorial agreement (PCA) was defined as all three replicates yielding the same categorical interpretation. If only two biological replicates of the MIC determination were valid, PEA and PCA were scored if both replicates were in agreement. Isolates with only one valid replicate were excluded from precision analyses. For each method, the proportion of isolates demonstrating PEA and PCA was calculated, and proportions were compared between the AD and PD methods using the Exact McNemar’s Test in RStudio. A *P* < 0.05 was considered statistically significant.

Statistical analysis was performed using indicated methods in R (R Core Team, Vienna, Austria) in RStudio (Posit Software, PBC, Boston, MA).

## Results

### Verification Results

The accuracy of fosfomycin PD compared with AD was evaluated against 78 isolates of species commonly isolated from urinary tract infections. Categorical and essential agreement are summarized in **Table 1**. An example of a PD plate is shown in **Fig. 1**.

**Figure 1.**
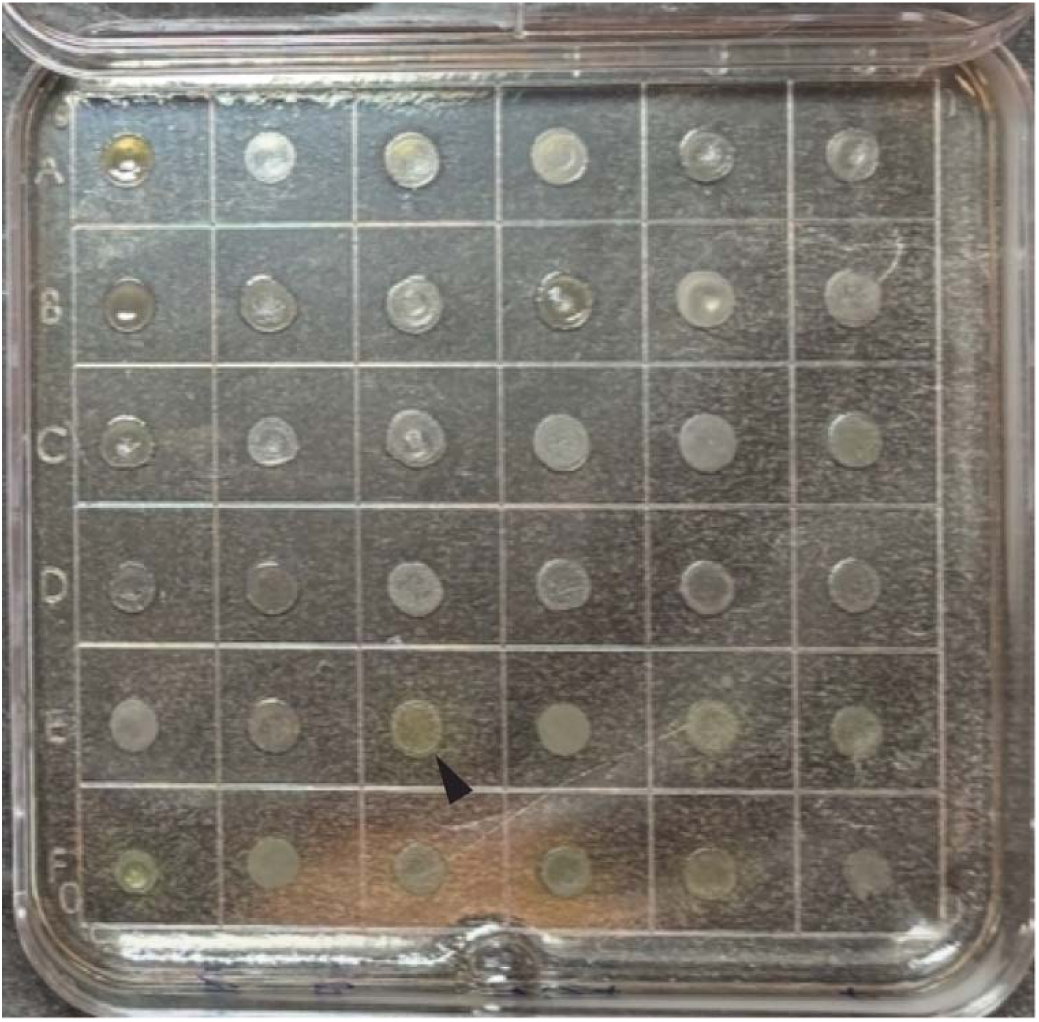
Poloxamer Dilution Testing Example. A representative poloxamer dilution plate inoculated with a Steer’s replicator is shown. Each spot of contained growth is a unique clinical isolate. Visibly apparent production of pyoverdine in *P. aeruginosa* isolates (arrowhead) can be observed on optically clear poloxamer media.

**Table 1.** Percent Accuracy of PD compared with Reference AD.

| Species | Total *n* | % CA (95% CI) | % VME (95% CI) | % ME (95% CI) | % MinE (95% CI) | % EA (95% CI) |
| --- | --- | --- | --- | --- | --- | --- |
| *E. coli* | 31 | 100.0 (89.0 – 100.0) | 0 (0.0 – 11.0) | 0 (0.0 – 11.0) | 0 (0.0 – 11.0) | 87.1 (71.1 - 94.9) |
| *Klebsiella* spp. | 20 | 60.0 (38.7 – 78.1) | 0 (0 – 16.1) | 10 (2.8 – 30.1) | 30.0 (14.5 – 51.9) | 55.0 (34.2 – 74.2) |
| *Other Enterobacterales* | 18 | 82.4 (59.0 - 93.8) | 0 (0 – 18.7) | 0 (0 – 18.7) | 17.7 (6.2 – 41) | 82.4 (59.0 – 93.8) |
| *P. aeruginosa* | 10 | 60.0 (31.2 - 83.1) | 0 (0 – 27.8) | 0 (0 – 27.8) | 40.0 (16.8 – 68.7) | 100.0 (77.2 – 100.0) |

Accuracy was compelling for *E. coli*, additional *Enterobacterales* species examined, and *Pseudomonas aeruginosa*. No VME were observed. Two major errors were observed for *K. pneumoniae. E. coli* strains, with the exception of the two pre-selected *fosA* strains from the AR-Bank (MIC > 512 µg/mL for both methods), were all highly susceptible, and essential agreement (EA) and categorical agreement (CA) were 87% and 100%, respectively (n=31). There was lower CA for *P. aeruginosa* and *Klebsiella* spp. based on the MICs of these species lying close to fosfomycin categorical breakpoints. EA was also lower for *Klebsiella* spp. based on several strains which tested at > 512 µg/mL for PD, and 256 or 128 µg/mL by AD. Overall, there was a bias towards higher and, therefore, more conservative PD MICs compared with AD, demonstrated in the scatter plot for all data (**Fig. 2**). Skipped dilutions, as has been previously observed for AD (5), occurred with AD and PD inconsistently among biological replicates. For *E. coli*, single and double skipped dilutions were approximately 6 times more frequent for AD than PD (**Table 2**, *P <* 0.0001 and *P* = 0.02, respectively, Fisher’s exact test). Proteus swarming was completely inhibited on PD at 16 to 20 hours of incubation (**Fig. 3**).

**Figure 2.**
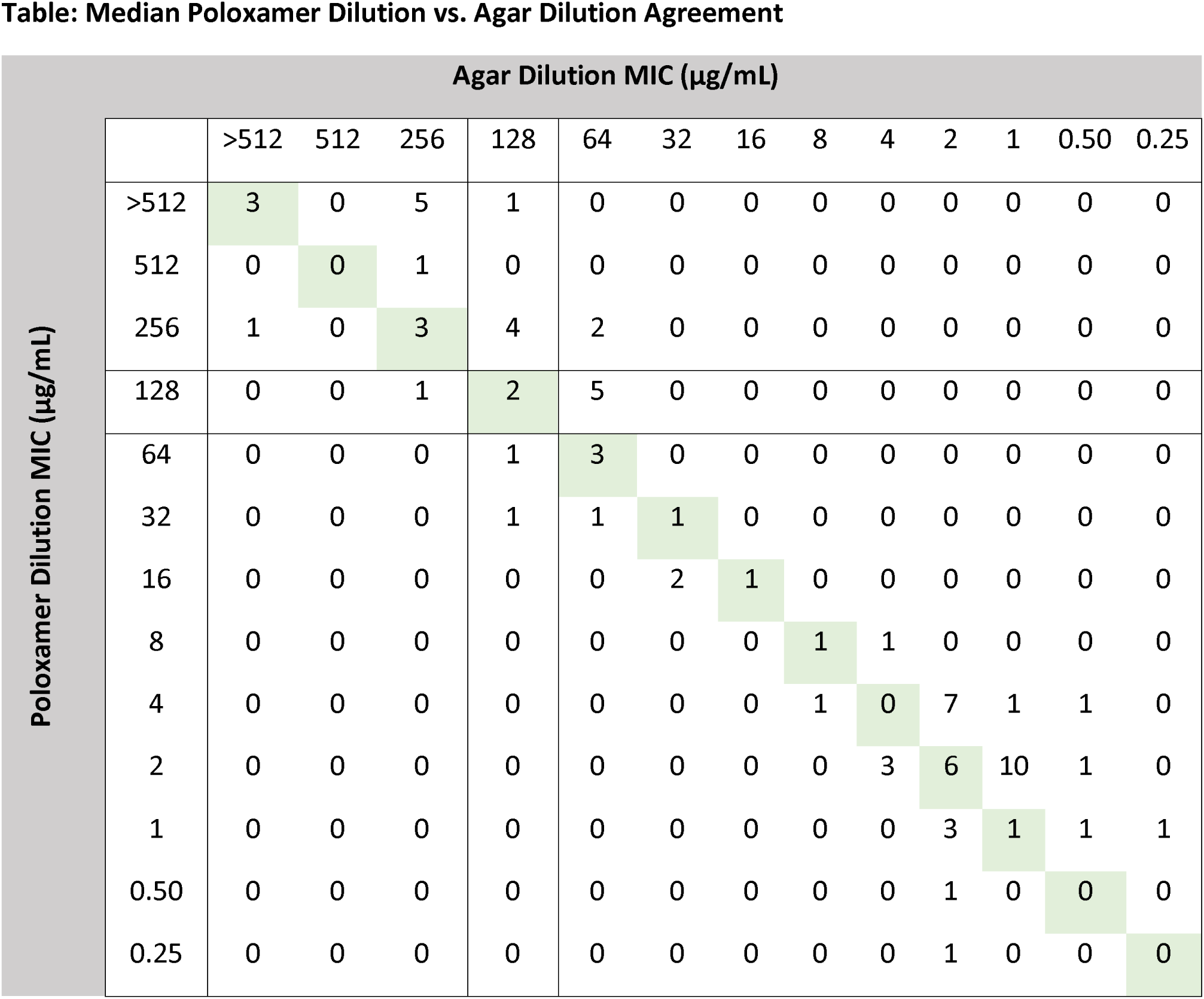
Scatterplot of PD versus AD MIC results. There was a slight positive bias (more conservative MIC call) for PD compared with AD. Per-species breakdown is provided in supplementary data (see **Table S2**). *E. coli* isolates ATCC 25922, AR-346, JEK-442 and AR-549 are not shown, but were all observed to be in EA and CA.

**Figure 3.**
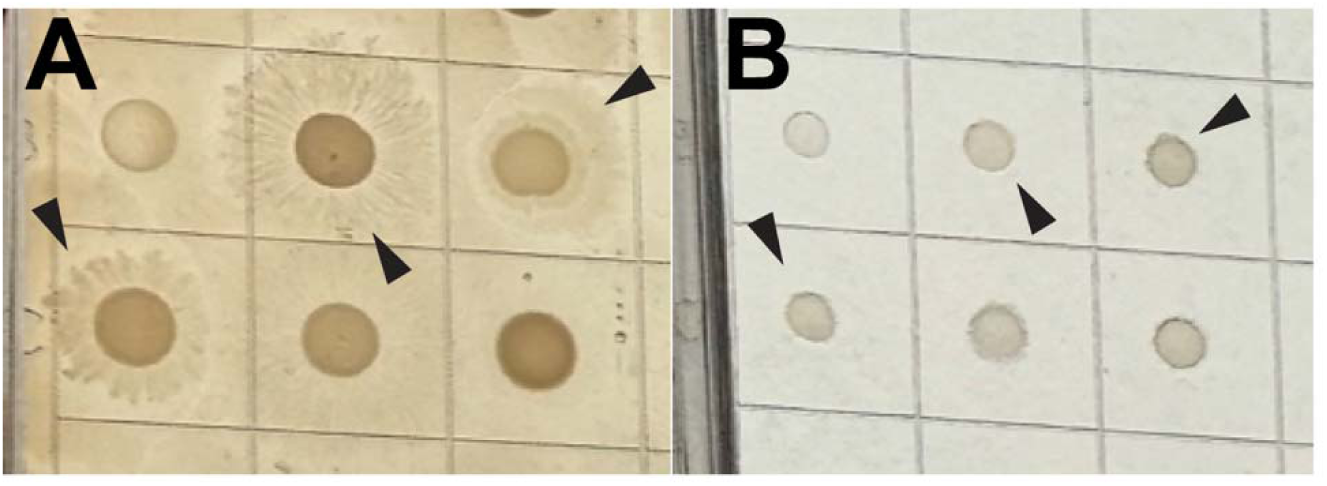
Differential Swarming Motility of *Proteus* spp. on Agar and Poloxamer. An image of bacterial isolate growth on an AD plate (A) and PD plate (B) both containing *Proteus* strains. Concentric swarming isolates of *Proteus* isolates on agar (arrowheads) are not observed on PD plates.

**Table 2.**
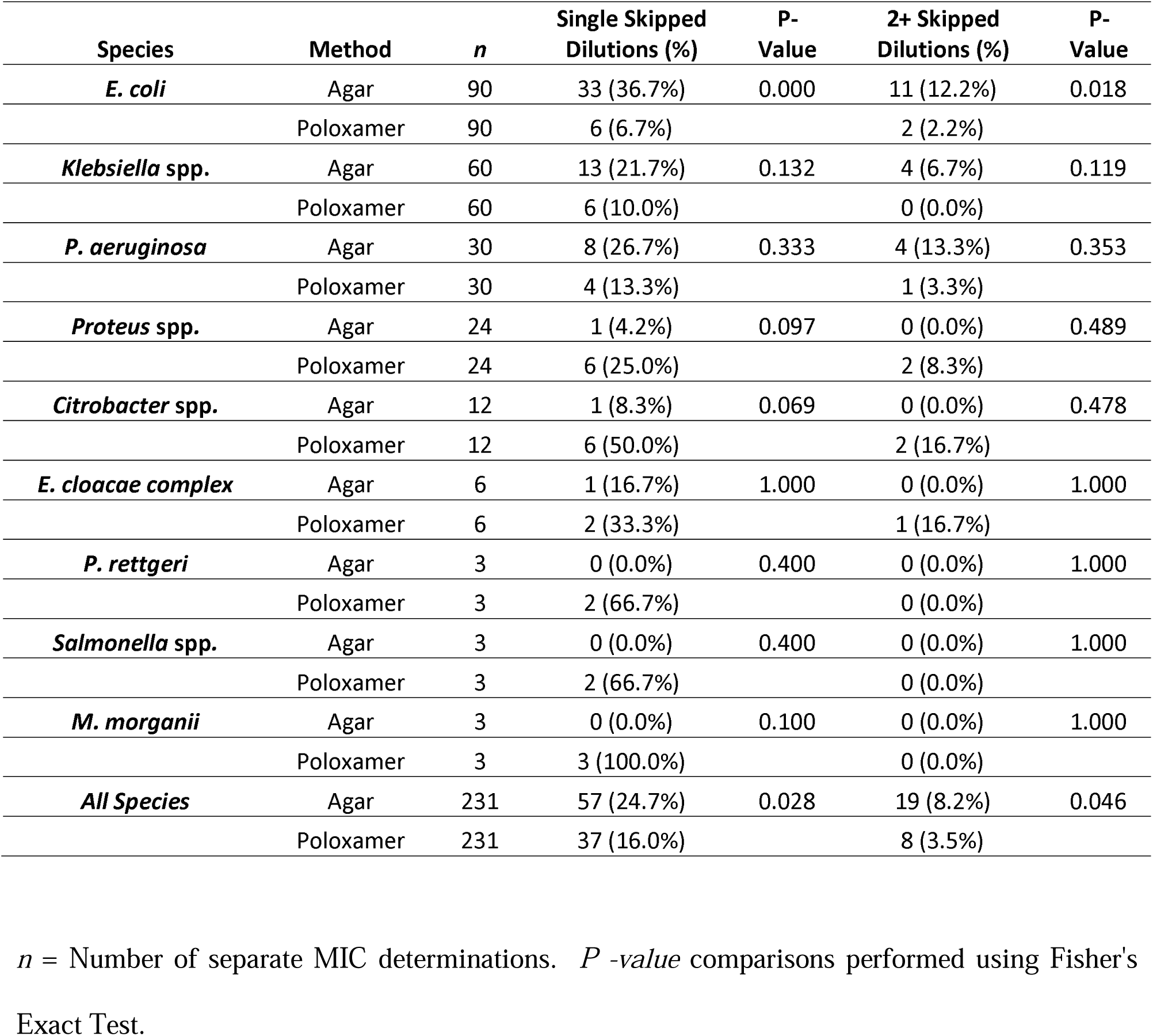
Percentage of single skipped dilutions observed in testing events for Poloxamer versus Agar Dilution.

### Precision

Precision agreement among the three biological triplicates for each isolate was calculated to compare reproducibility of PD compared with AD. Interestingly, precision essential agreement for PD (74%) was higher than for AD (63%), however, this did not reach statistical significance (P = 0.2). The lower PEA for AD was attributable in part to increased variability in measured results caused by skipped dilutions. Precision categorical agreement (PCA) for PD (81.6%) was marginally higher than AD (76.3%), a difference that likewise did not reach statistical significance (*P* = 0.5, Exact McNemar Test, **Table S3**).

## Discussion

In this study, we highlight the potential of poloxamer dilution (PD) as an accurate and precise, on-demand alternative to agar dilution (AD) for antimicrobial susceptibility testing. Using fosfomycin testing as a use case, PD performed well relative to the AD reference method whose performance was compromised by the more frequent occurrence of skipped dilutions (5). For *E. coli*, PD clearly distinguished susceptible from resistant *fosA* strains, and PD also had reasonable essential agreement for *Enterobacterales* and *P. aeruginosa* compared with AD. Errors were almost always conservative across species. *Klebsiella* spp., which constituted approximately 25% of isolates tested, most of which were fosfomycin intermediate or resistant by AD, contributed to approximately 50% of the total categorical and essential agreement errors, all of which were conservative (i.e. reading at a higher MIC by PD). Therefore, demonstration of fosfomycin susceptibility by PD could be used broadly to confirm that urinary drug concentrations are likely to reach levels sufficient for organism clearance, while acknowledging that PK/PD data remain insufficient for organisms other than *E. coli* and *E. faecalis*.

Because of its thermoreversible properties, autoclaved poloxamer media can be aliquoted and stored sterile in liquid form at 4 °C, allowing immediate setup of PD testing. This same property permits antibiotics to be added to cold, liquified poloxamer without risking heat-associated degradation that can occur during preparation of agar dilution plates.

PD testing can also be performed in microwell plates (**Fig. S1**). In previous work, we validated a digital dispensing method (DDM) using the HP D300 inkjet printer, which enables rapid, on-demand generation of custom antimicrobial susceptibility panels for clinical laboratory testing (6, 8, 13–15). The printer dispenses precisely sized droplets of antibiotic stock solutions (“ink”) to achieve any desired final concentration within each micro- or macro-dilution well. Using this approach, reference-method–equivalent doubling dilution MIC panels can be produced within minutes. In an example of micro-well PD testing, digital dispensing was used to print the appropriate doubling dilutions of fosfomycin stock solution into a dry 48-well plate. Previously prepared, sterile, liquid Mueller–Hinton–Poloxamer (MH-P) medium containing glucose-6-phosphate was then dispensed into the wells using a Combitip positive displacement repeat pipetter. After gentle mixing, plates were allowed to solidify at room temperature, and testing was piloted with 25 *E. coli* clinical isolates using CE-marked Liofilchem pre-made 24-well agar dilution plates as the reference comparator. Categorical agreement (CA) was 96% with one minor error, and evaluable essential agreement (EA) was also 96% (**see Table S6**), supporting the suitability of the microwell PD testing format.

Taken together, these findings position PD as a practical and reliable method that can be implemented within the local clinical microbiology laboratory, bringing reference-level antimicrobial susceptibility testing capabilities closer to the site of patient care. By enabling flexible, on-demand preparation of high-quality dilution panels, PD technology has the potential to expand testing portfolios and substantially reduce turnaround time while maintaining the analytical rigor of reference methods. Further studies are warranted to explore the applicability of this approach across a broader range of bacterial species and antimicrobial classes.

## Acknowledgements

This work was supported by a Novel Therapeutics Delivery Grant from Massachusetts Life Science Center to J.E.K. M.T.J.U. received a Carole Shapazian Research Co-op Fellowship to support a research co-op experience from Northeastern University. The HP D300 digital dispenser used in these studies was provided by TECAN (Morrisville, NC). TECAN had no role in study design, data collection, or interpretation.

