## Supplementary material for "Poloxamer dilution as an on-demand alternative to agar dilution-based antimicrobial susceptibility testing": S1 Figure

**Figure S1. Poloxamer dilution testing in 48-well microplate format.** A microwell format allows testing of a full doubling-dilution range for multiple isolates, including a QC strain, in a single plate.

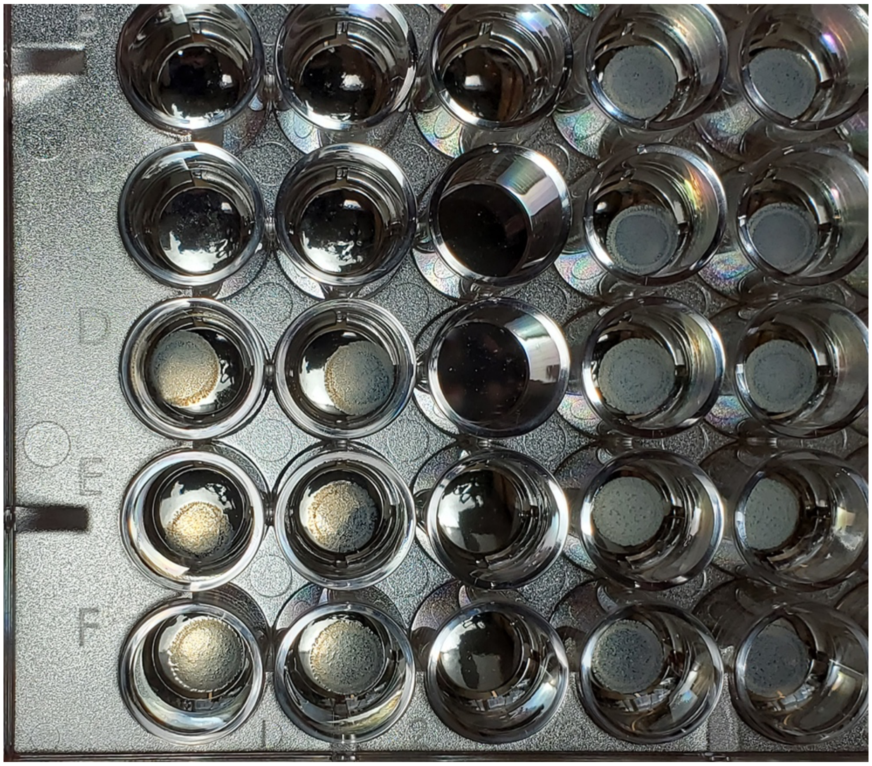
